# Cytotoxicity, Mutagenicity and Genotoxicity of Electronic Cigarettes Emission Aerosols Compared to Cigarette Smoke: the REPLICA project

**DOI:** 10.1101/2022.10.28.514205

**Authors:** Rosalia Emma, Virginia Fuochi, Alfio Distefano, Sonja Rust, Fahad Zadjali, Mohammed Al Tobi, Razan Zadjali, Zaina Alharthi, Roberta Pulvirenti, Pio Maria Furneri, Riccardo Polosa, Massimo Caruso, Giovanni Li Volti

## Abstract

During the last decade electronic cigarettes (e-cigarettes) have been studied as an alternative devices to the tobacco cigarette, but with better safety for the health of smokers, so as to create a new approach to smoking addiction, such as the “smoking harm reduction”. This new approach, suggested by a part of the scientific world, aroused interest and debates in the regulatory field, involving all the major regulatory bodies and often creating divergences from nation to nation on the rules driving the production, distribution and consumption of these alternative products. Many studies have been conducted both in vitro and in vivo, to clarify the effects of the e-cigarette compared to the classic one. In this context, the Center of Excellence for the Acceleration of HArm Reduction (CoEHAR) was established within the University of Catania (Italy) and the multi-center project, created under its leadership, the REPLICA project, which aims to replicate in vitro studies originally conducted by tobacco and e-cigarette manufacturers, in order to verify the robustness and replicability of the data. In this work the REPLICA Team replicated part of the work published by Rudd and colleagues in 2020, which aims to establish the aerosol-induced cytotoxicity, mutagenesis and genotoxicity of a pod system e-cigarette aerosol compared to tobacco cigarette smoke. As in the original paper, we performed Neutral Red Test (NRU) for the evaluation of cytotoxicity, AMES test for the evaluation of mutagenesis and In Vitro Micronuclei (IVM) assay for the evaluation of genotoxicity on cells treated with cigarette smoke or e-cigarette aerosol. The results obtained showed high cytotoxicity, mutagenicity and genotoxicity induced by cigarette smoke, but slight or no cytotoxic, mutagenic and genotoxic effects induced by the e-cigarette aerosol. The data obtained support those previously presented by Rudd and colleagues, although we have highlighted some methodological flaws of their work. Overall, we can affirm that the results obtained by Rudd and colleagues have been established and our data also confirm the idea that e-cigarette aerosol is much safer and less harmful than cigarette smoking, making it a useful device in smoking harm reduction.

## Introduction

In recent years, electronic cigarettes (e-cigarettes) have been gaining popularity as a safer alternative to tobacco products. A recent safety review by the Committee on Toxicity of Chemicals in Food, Consumer Products and the Environment (COT) of the Public Health England stated that the risk of adverse health effects from vaping products, produced according to appropriate manufacturing standards and used as recommended, is much lower than from tobacco cigarettes. However, the possible health risk of inhalation of flavour and their thermally-derived products is still “an area of uncertainty” [1].

The interest in public health and in regulatory policies regarding toxicological aspect of vapor products has increased worldwide. This was mainly due to reported cases of lung injury (EVALI) associated to improper use of e-cigarettes for tetrahydrocannabinol (THC) consumption as well as vitamin E additives [2]. Some countries banned the e-cigarette, such as India, Australia, Oman, Egypt, Colombia, etc. Whereas, in other countries the government institutions have imposed rules for the marketing of e-cigarettes and e-liquids. The U.S. Food and Drug Administration (FDA) published requirements and industry guidance in order to regulate the premarket tobacco product applications for electronic nicotine delivery systems [3].Also, the European Union Tobacco Products Directive (2014/40/EU) provided rules on e-cigarettes premarket requirements [4]. Both FDA and UE directives required to manufacturers and importers of vape products to submit ingredient, emissions and toxicological data to authorities for the marketing of their products [3, 4].

For the assessment of toxicological potential of e-cigarettes, international guidelines (the International Conference on Harmonisation S2(R1) (2011), the UK Committee on Mutagenicity of Chemicals in Food, Consumer Products and the Environment (2011), Health Canada (2005) and the Cooperation Centre for Scientific Research Relative to Tobacco (CORESTA) (2004)) recommend the use of a battery of *in vitro* tests as pre-clinical assessment strategy. Particularly, these guidelines require the evaluation of various toxicological endpoints through the use of multiple assays, such as the bacterial reverse mutation (AMES) assay for mutagenicity, *in vitro* micronucleus (IVM) for genotoxicity, combined with neutral red uptake (NRU) for acute cytotoxicity evaluation [5, 6]. These three *in vitro* toxicity tests are widely used as standard assays to assess the toxicity of tobacco products and e-cigarettes [7].

The main e-cigarette manufacturers, including tobacco companies, published studies on their product evaluations, including emissions, cytotoxicity, genotoxicity and mutagenicity data [8-11]. Independent replication of these studies can provide verification of the company’s findings to establish the credibility of the data and support regulation of electronic cigarettes. Faulty results are ill-informing policies and have detrimental effects on research practices undermining public trust in science and eventually affecting health and social care practices. The multicenter REPLICA project was created to replicate the high-profile studies coming from the R&D of Tobacco Companies in order to assess validity of the original work put under scrutiny (https://replica.coehar.org/).

In the last phase of REPLICA project, the Italian team (CoEHAR, University of Catania – LAB-A) and the partner in Oman (Sultan Qaboos University – LAB-B) conducted a replication study of a paper published by Rudd and colleagues from the Imperial Brands PLC [8]. In summary, this paper reports comparative data for aerosol emissions and in vitro toxicity, using the neutral red uptake (NRU), the bacterial reverse mutation (AMES), and in vitro micronucleus (IVM) assays, for a pod-system e-cigarette (*my*blu™) compared with 3R4F reference cigarette smoke. They observed that many of the harmful and potentially harmful constituents found in cigarette smoke were not detected in e-cigarette aerosol. Using established *in vitro* biological tests, e-cigarette aerosol did not display any mutagenic or genotoxic activity under the conditions of test. By contrast, 3R4F cigarette smoke displayed mutagenic and genotoxic activity. E-cigarette aerosol was also found to be 300-fold less cytotoxic than cigarette smoke in the neutral red uptake assay.

Here we replicated the *in vitro* biological test to investigate on cytotoxic, mutagenic and genotoxic activity of *my*blu™ e-cigarette aerosol versus 1R6F cigarette smoke using similar methods in order to assess the results obtained by Rudd and colleagues [8]

## Methods

### Test products

Unlike Rudd and colleagues, we used the 1R6F reference cigarette (University of Kentucky, Center for Tobacco Reference Products, Lexington, KY, USA), since the 3R4F are no longer produced by the University of Kentucky. 1R6F and 3R4F reference cigarettes are very similar and only slight differences were reported regarding smoke chemistry and *in vitro* assays [12]. Prior to every experimental session, the 1R6F cigarette were conditioned for a minimum of 48 h at 22 ± 1 °C and 60 ± 3% relative humidity, according to ISO 3402:1999 [13].

The same electronic cigarette used by Rudd and colleagues, *my*blu™ (Imperial Brands PLC, Bristol, United Kingdom), was used for this replication study. The *my*blu™ is aa “closed pod-system” e-cigarette consisting of two elements: a rechargeable battery (battery capacity, 350 mAh) and a replaceable e-liquid pod with 1.5 mL volume and a coil resistance of 1.3 Ω (cartomizer). The tobacco flavored e-liquids with 1.6% (w/w) nicotine were used for the experiments. All the *my*blu™ e-cigarettes and *my*blu pods were purchased from Italian retailers.

### Smoke and vapour exposure

The selected products were tested on standardized equipment simulating smoking topography: the LM1 Smoking Machine (Borgwaldt, Hamburg, Germany) was used to smoke the 1R6F cigarettes following the Health Canada Intense (HCI) regime (puff volume, duration and frequency of 55 mL, 2 s and 30 s (55/2/30), with bell shaped profile and hole vents blocked), accredited under ISO/TR 19478-2:2015.

The LM1 is a direct exposure system and does not perform smoke dilution unlike the smoking machine used by Rudd and colleagues. The LM4E Vaping Machine (Borgwaldt, Hamburg, Germany) was used to vape *my*blu™ following the “CORESTA Reference Method n. 81” (CRM81) regimen (55 ml puff volume, drawn over 3 s, once every 30 s with square shaped profile), accredited into ISO 20768:2018.

An air-liquid exposure system different from the original paper by Rudd et al. was used to expose BEAS-2B to perform NRU assay and V79 cells to perform IVM assay. The BAT (British American Tobacco) aerosol exposure chambers were used to expose *in vitro* cells at the air-liquid interface (ALI) to cigarette smoke and aerosols from e-cigarettes. In particular, cells cultured in Transwell inserts were deprived of the apical medium and placed in the exposure chambers containing the medium in the lowest compartment, keeping wet the basal face of the cells, and then connected to the smoking or vaping machine to deliver undiluted whole smoke or whole aerosol to the apical face of cells (Figure 1) [14].

**Figure 1.**
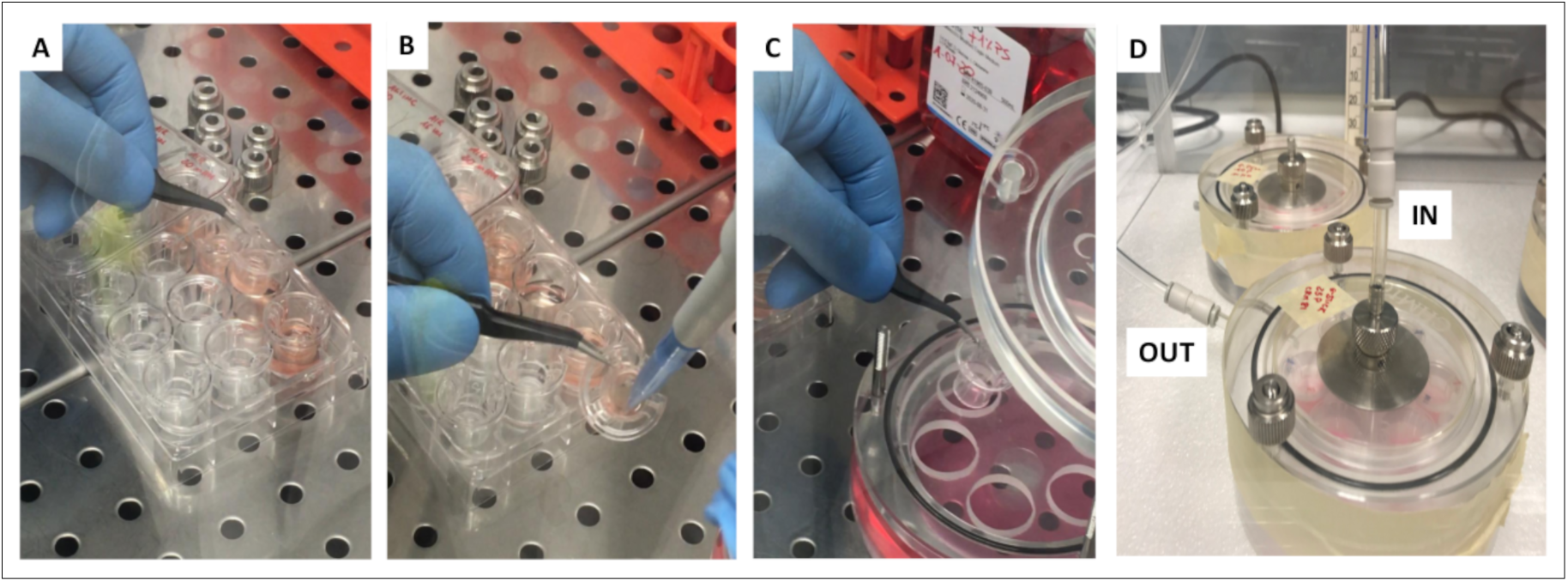
Air-liquid interface exposure system

For the cytotoxicity evaluation of BEAS-2B, 1R6F cigarette smoke was delivered undiluted from 1 to 8 puffs. The *my*blu aerosol was delivered undiluted from 20 to 140 puffs, as reported by Rudd et al. Cytotoxicity evaluation was also performed to establish the EC50 of V79 cells exposed to 1R6F undiluted smoke prior to the IVM assay. In that case, V79 cells were exposed to 1R6F smoke from 2 to 30 puffs. Based on these results, we performed the 1R6F smoke exposure for the IVM delivering from 1 to 4 puffs. The *my*blu aerosol was delivered undiluted from 20 to 100 puffs for the IVM assay, as reported by Rudd et al.

For the AMES assay, 1R6F cigarette smoke and the relative AIR control, or *my*blu aerosol and the relative AIR control were delivered to the bacterial suspensions contained into their corresponding impingers at room temperature under protection from direct light. A flushing step with filtered ambient air was applied after each puff. The number of puffs are reported in table 1 for the 1R6F exposure and in table 2 for the *my*blu exposure. In particular, the impinger INLET for the 1R6F smoke exposure was connected to LM1 smoking machine (1R6F smoke) and to a LM4E channel equipped with a 44 mm CFP filter pad (AIR interpuff) by means of a double one-way valve, whereas the impinger INLET for the *my*blu e-cigarette aerosol exposure was connected to two different LM4E channels, one of which equipped with e-cigarette and the other with a 44 mm CFP filter pad (AIR interpuff).

**Table 1.** Exposure conditions for the bubbling of bacterial suspension (AMES assay)

| Conditions | Puff number<br>(LM1) | AIR interpuff number<br>(LM4E) | Conditions | Puff number<br>(LM4E port 1) | AIR interpuff number<br>(LM4E port 2) |
| --- | --- | --- | --- | --- | --- |
| **1R6F (HCI regime)** |  |  | ***myblu* e-cig (CRM81 regime)** |  |  |
| 1 cig | 9 | 9 | 60 puffs | 60 | 60 |
| 2 cigs | 18 | 18 | 120 puffs | 120 | 120 |
| 3 cigs | 27 | 27 | 180 puffs | 180 | 180 |
| 4 cigs | 36 | 36 | 240 puffs | 240 | 240 |
| 5 cigs | 45 | 45 | 300 puffs | 300 | 300 |
| **AIR Control (1R6F) – HCI** |  |  | **AIR Control (*myblu* e-cig) – CRM81** |  |  |
| 1 cig | 9 | / | 60 puffs | 60 | / |
| 2 cigs | 18 | / | 120 puffs | 120 | / |
| 3 cigs | 27 | / | 180 puffs | 180 | / |
| 4 cigs | 36 | / | 240 puffs | 240 | / |
| 5 cigs | 45 | / | 300 puffs | 300 | / |

**Table 2.**
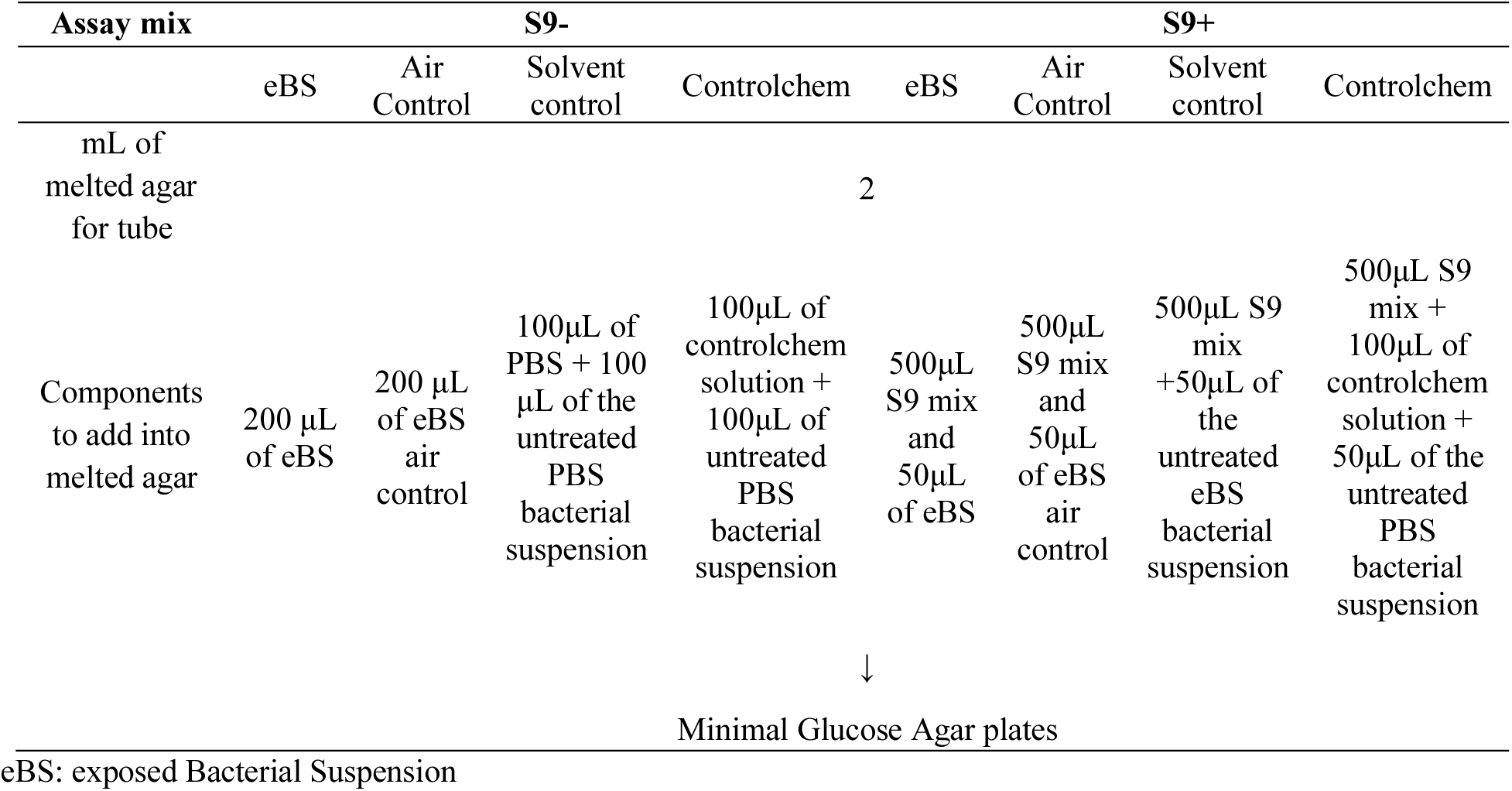
Schematic representation of the assay. The example is intended for one bacterial strain only and for both S9- and S9+ mix.

### Cell cultures

Normal bronchial epithelial cells (BEAS-2B / ATCC-CRL-9609) were cultured in collagen coated flasks using the bronchial epithelial growth medium supplemented with Lonza Bullet Kit (BEGM, Lonza CC-3170), as described by ATCC culture instructions. Hamster lung fibroblast cells (V79 / ICLC-AL99002) were cultured using Dulbecco’s Modified Eagle Medium-high glucose (DMEM-hg, Thermo Fisher Scientific) with 10% FBS, 2 mM L-Glutamine, 50 U/mL penicillin, and 50 μg/mL streptomycin, as described by ICLC instructions.

### Cytotoxicity evaluation: NRU assay

Cytotoxicity evaluation was performed using the BEAS-2B cells by using the NRU assay [15, 16]. Moreover, cytotoxicity evaluation was performed with 1R6F whole smoke for the V79 prior to genotoxicity evaluation in order to establish the puff number conditions to be used.

Prior to exposure, 300 μl of BEAS-2B cell suspension (BEGM supplemented with Lonza Bullet Kit and with 20mM of HEPES buffer) was seeded in 24-well Transwell® inserts at density of 150.000 cells/well, and incubated for 20±3 hours. After incubation, the apical cell culture medium was removed, and the Transwell® inserts were transferred into the corresponding exposure chamber filled with 25 ml of DMEM-hg supplemented with 50 U/mL penicillin and 50 μg/mL streptomycin in order to proceed to the smoke/vapor ALI exposure. After ALI exposure, each insert was transferred in a new 24-well plate filled with 500 μl and 300 μl of fresh BEGM (supplemented with Lonza Bullet Kit + 20mM of HEPES buffer) respectively at the basal and apical compartments. Next, the cells were incubated for a recovery period of 65±2h. The day before NRU assay, the NRU solution was prepared in BEGM medium at ratio 1:65 (0.05 g/L) plus HEPES buffer at 20 mM and placed in incubator at 37 °C 5% CO_2_. The day of NRU assay, the NRU solution was filtered prior to use. The culture medium was removed from the apical and basal compartments of each culture insert. The cells were washed twice with pre-warmed PBS, then incubated with neutral red solution (500 μl at the bottom and 300 μl at the top) for 3 h at 37°C in 5% CO_2_ and a humidified atmosphere. After incubation, cells were washed twice with pre-warmed PBS to remove unincorporated dye. The incorporated solution was eluted from the cells by adding 330 μl of destain solution (50% ethanol, 49% distilled water, 1% glacial acetic acid v:v:v) to each insert, and incubated for 10 min at 300 rpm on a plate shaker. Extracts were transferred to a 96-well plate in duplicate (100 μl per well) and optical density of neutral red extracts was read with a microplate spectrophotometer at 540 nm using a reference filter of 630 nm. Blank inserts (without cells) were used to assess how much neutral red solution stained the Transwell® membranes and the mean of background values was subtracted from each measurement. The same procedure was used for the cytotoxicity evaluation of V79 cells. In brief, 300 μl of V79 cell suspension (DMEM-hg supplemented with 10% FBS, 2 mM L-Glutamine, 50 U/mL penicillin, 50 μg/mL streptomycin, and 20 mM HEPES) was added in 24-well Transwell inserts at density of 100.000 cells/well, and incubated for 24 hours. The day of ALI exposure, the Transwell® inserts (without apical cell culture medium) were transferred into the corresponding exposure chamber filled with 25 ml of DMEM-hg supplemented with 50 U/mL penicillin and 50 μg/mL streptomycin. After ALI exposure, each insert was transferred in a new 24-well plate filled with 500 μl and 300 μl of fresh DMEM-hg (supplemented with 10% FBS, 2 mM L-Glutamine, 50 U/mL penicillin, 50 μg/mL streptomycin, and 20 mM HEPES) respectively at the basal and apical compartments. Next, the cells were incubated for a recovery period of 24 hours. The NRU solution was prepared in DMEM-hg at ratio 1:65 (0.05 g/L) plus HEPES buffer at 20 mM. The next steps of NRU assay for V79 cells were the same of BEAS-2B and are described above.

### Mutagenicity evaluation: AMES assay

The *in vitro* mutagenicity of fresh 1R6F smoke and *myblu* aerosols was determined using Ames test [17] as described by Rudd et al. [8] with some modifications, and it was conducted only by LAB-A. The Ames screen was employed using *S. typhimurium* TA98 and TA100 strains (Trinova Biochem GmbH) ±S9 treatment, conducted in accordance with OECD (Organization for Economic Cooperation and Development) test guideline 471 [18].

Briefly, bacterial cultures of the TA98 and TA100 strains were prepared in 25 mL Nutrient Broth No.2 (OXOID) by inoculating one bacterium-coated CRYO-glass bead followed by incubation overnight at 37°C with shaking at 120 rpm. Then, bacterial suspensions were prepared by centrifugation of 25 mL cultures at 1800 g for 20 minutes at 4°C, and the pellet was resuspended in 12 mL of Ca^2+^, Mg^2+^-free Dulbecco’s phosphate buffered saline (PBS). 10 mL of the bacterial suspensions were placed in the corresponding impingers and exposed to test aerosols/smoke as described above (exposed Bacterial Suspension or eBS). For each experiment an aliquot of PBS with untreated PBS bacterial suspension (for the assay without S9 mix) and S9 mix with untreated PBS bacterial suspension (for the assay with S9 mix) as internal negative controls (Solvent control).

After each exposure, the bacterial suspensions were immediately used for AMES screening by following manufacturer’s protocol (Salmonella Mutagenicity Test Kit, MOLTOX^®^). Briefly, an aliquot of bacterial suspensions and relative reagents were added to sterile 15 mL test tubes as described in Table 2. The solution was thoroughly mixed and then decanted on top of a minimal glucose agar plate, covered and set aside to solidify. When the top agar was solidified, the plates were inverted and placed in an incubator at 37°C. After 48-72 hours of incubation, the number of revertant colonies growing on the plates was counted manually.

Controlchem™ Mutagens were used as positive controls for both *S. typhimurium* stains TA98 and TA100 (see Table 3). Each concentration of test aerosols or smoke and positive controls was tested in triplicate.

**Table 3.**
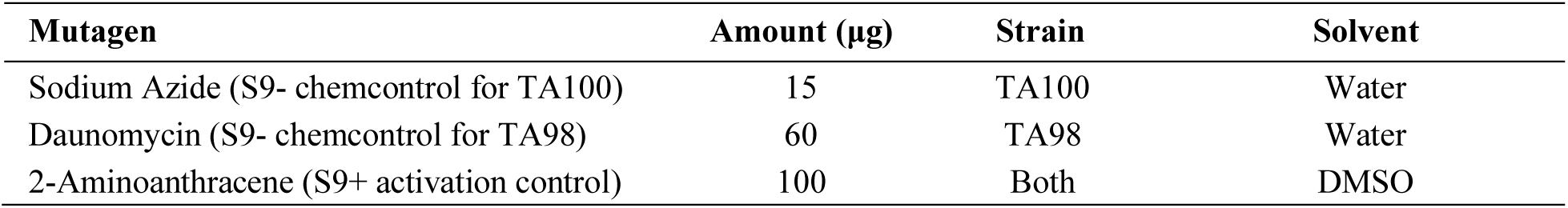
Controlchem™ Mutagens and Related Solvents.

| Mutagen | Amount (µg) | Strain | Solvent |
| --- | --- | --- | --- |
| Sodium Azide (S9- chemcontrol for TA100) | 15 | TA100 | Water |
| Daunomycin (S9- chemcontrol for TA98) | 60 | TA98 | Water |
| 2-Aminoanthracene (S9+ activation control) | 100 | Both | DMSO |

### Genotoxicity evaluation: IVM assay

The IVM assay was performed in concordance with OECD test guideline no. 487 [19] with and without

S9 metabolic activation, and it was conducted only by LAB-A. Genotoxicity evaluation of whole fresh tobacco smoke and e-cigarette aerosol was performed using the hamster lung V79 cell line (ICLC-AL99002). The day before cell exposure, V79 cells were seeded in 24-well Transwell® inserts (0.4 μm pore membrane) at density of 10*10^4^ with 200 μl of DMEM-hg supplemented with 10% of FBS. The inner wells of each 24-well plate were filled with 500 μl of DMEM-hg supplemented with 10% of FBS. The V79 cells were incubated for 24 hours at 37 °C and 5% CO_2_. After 24h of incubation, the apical medium was removed, and the inserts were transferred into the exposure chamber filled with 25 ml of DMEM-hg with the addiction of HEPES buffer (20 mM final concentration) in the basal compartment. The exposure to 1R6F whole smoke and myblu e-cigarette whole aerosol are described in the previous section on “<u>Smoke and vapour exposure”.</u> After the exposure, each insert was transferred in a new 24-well plate filled with 500 μl of DMEM-hg supplemented with HEPES buffer (20mM). For the IVM with S9, each insert containing exposed V79 cells, was filled with 300 μl of S9 mix at 10% in the apical compartment, and then incubated for 3 hours at 37 °C. After incubation, the apical S9 medium was removed, the V79 cells were covered with DMEM-hg supplemented with HEPES buffer (20mM), and incubated for 24 hours to allow for at least one cell division cycle. For the IVM without S9, inserts with exposed V79 cells were filled with DMEM-hg supplemented with HEPES buffer (20mM) and incubated for 24 hours at 37 °C and 5% CO_2_. Positive controls, including cyclophosphamide A (CAS 6055-19-2) for the S9 fraction and mitomycin C for the IVM without S9, were used. All the tested conditions were assessed in triplicates. After 24 hours of recovery, the cells were detached and counted using the Guava^®^ Muse^®^ Cell Analyzer using the Muse^®^ Count & Viability Kit (Luminex Corp.). The V79 cells were then seeded in 96-well plate (CellCarrier Ultra-96 Black, Optically Clear Bottom - PerkinElmer) at density of 10*10^3^ per well and incubated for 24h. Next, the cells were fixed with 4% (PFA) (paraformaldehyde) for 20 minutes at room temperature. After fixation, the cells were washed once with PBS and then stained with DAPI (1 lg/mL). Micronuclei assessment was performed by using Harmony High-Content Imaging and Analysis Software.

### Statistics

All raw data were collected and processed using Excel software (Microsoft, Redmond, WA, USA). For the cytotoxicity evaluation (NRU), data were expressed as percentage to the AIR control. The EC50 values for each exposure (1R6F and *my*blu) were calculated by fitting a sigmoidal dose-response curve with a variable slope to determine the best fit values for the 1R6F log EC50 of 7 parameter nonlinear regression model and for the *my*blu log EC50 of 7 parameter nonlinear regression model. Moreover, comparison of *my*blu results and AIR control was carried out by ANOVA followed by Dunnett’s post hoc multiple comparison test.

Data from AMES assay (mutagenicity evaluation) were reported as fold change relative to AIR control [20], and calculated as follow:

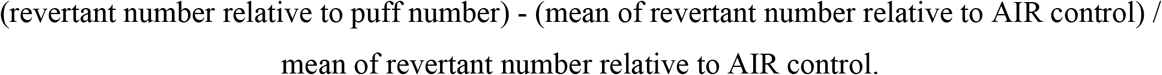

Linear regression analyses for each strain were performed to evaluate the mutagenic activity. Moreover, comparisons among 1R6F and respective controls and among *my*blu and respective controls were performed by using mixed-effect model or ANOVA followed by Tukey’s post hoc multiple comparison test.

Genotoxicity data were analyzed by linear regression of 1R6F or *my*blu dose-response slopes with comparison between slopes. Comparisons among the different dose of smoke or aerosol and respective controls were performed by ANOVA followed by Tukey’s (IVM without S9 activation) or Dunnett’s (IVM with S9 activation) post hoc multiple comparison tests.

All analyses were considered significant with a p value < 0.05. GraphPad Prism 8 software was used for data analysis and generation of graphs.

## Results

### Cytotoxicity: effect of cigarette whole smoke and *my*blu whole aerosol on cell viability

After exposure with whole smoke from 1R6F reference cigarettes, BEAS-2B cell viability drastically decreased as early as 2 puffs until complete cell death at 4 puffs with an EC50 value of 1.71 puffs (Figure 2A). Instead, unlike Rudd and colleagues, the EC50 value could not be calculated for *my*blu exposure due to low cytotoxity at 140 puffs (Figure 2B). We observed a reduced cell viability starting from 80 puffs to 140 puffs, which did not decrease below the 80% of viability. Particularly, significant decreased cell viability was observed for 80 (p= 0.003), 100 (p= 0.008), and 140 puffs (p= 0.002) compared to the AIR control.

**Figure 2.**
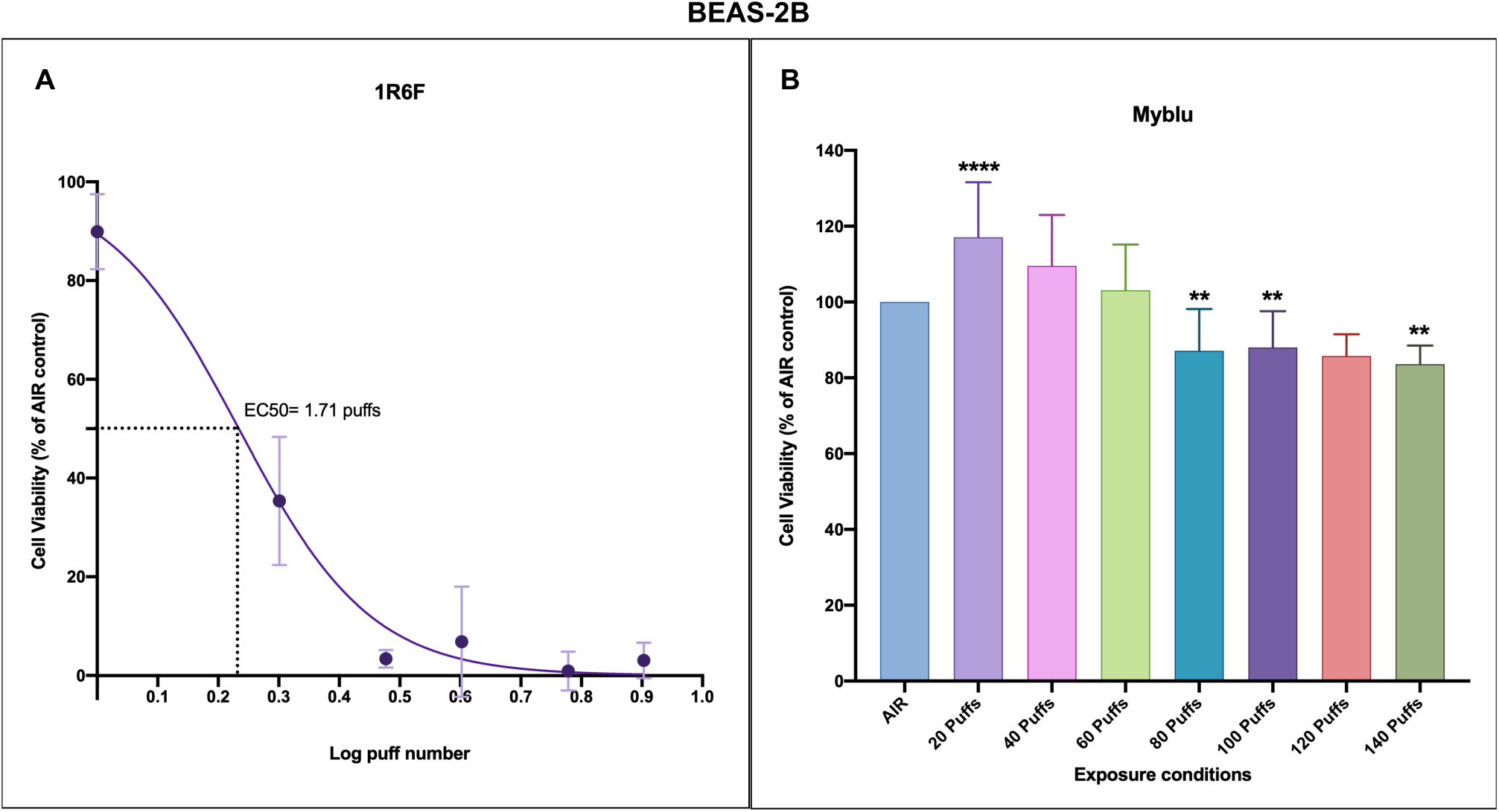
Cytotoxicity evaluation of BEAS-2B cells after exposure to 1R6F cigarette smoke and *my*blu aerosol. (**A**) Dose response curve of BEAS-2B cells exposed to 1R6F smoke showed an EC50 value of 1.71 puffs (Log EC50= 0.23 puff). (**B**) Barplot representing the BEAS-2B cell viability results after exposure to *my*blu aerosol. All data are reported as percentage of the respective AIR control, and displayed as mean ± standard deviation (SD). **p< 0.01; ****p< 0.0001.

In addition to Rudd and colleagues, we observed microscopically the cells exposed to both 1R6F smoke and *my*blu aerosol at 24, 48, and 65 hours. Exposure of BEAS-2B cells to 1R6F smoke induced morphological changes of cells with alterations in cell volume, nucleus volume, and cell sphericity at all the time-points. Similar morphological changes of BEAS-2B cells were observed after exposure to *my*blu aerosol starting from 80 puffs at 24 h. But, a reversal of morphological changes was observed from 48 h to complete recovery at 65 h.

### Mutagenicity effect of 1R6F cigarette smoke and *my*blu aerosol

The negative controls (Solvent control) were in the normal range based on our laboratory experience and to literature data [20]. Also, the positive controls (i.e., Chem Controls; Sodium Azide, Daunomicyn, and 2-Aminoanthracene) were in the range reported in the manufacturing manual (Trinova Biochem GmbH – Germany). No significant differences were observed between Solvent controls and AIR controls for all the tested conditions. Instead, significant differences were observed between Solvent controls or AIR controls and the respective Chem Controls (p< 0.0001).

Exposure to 1R6F cigarette smoke induced significant increase in revertants in a dose-dependent manner for both TA98 (up to nearly 4-fold change to AIR control; p< 0.0001) and TA100 (up to 1-fold change to AIR control; p= 0.002) with S9 metabolic activation. No significant increase in revertants from zero was observed for TA98 strain with S9 metabolic activation after exposure to *my*blu aerosol. Instead, a slight increase of revertants for TA100 S9+ (up to approximately 0.2-fold change to AIR control; p= 0.005) was observed after *my*blu aerosol exposure (Fig. 3). The linear regression results of mutagenic activity in *Salmonella tiphimurium* (TA98 and TA100) with S9 metabolic activation are reported in table 4.

**Table 4.**
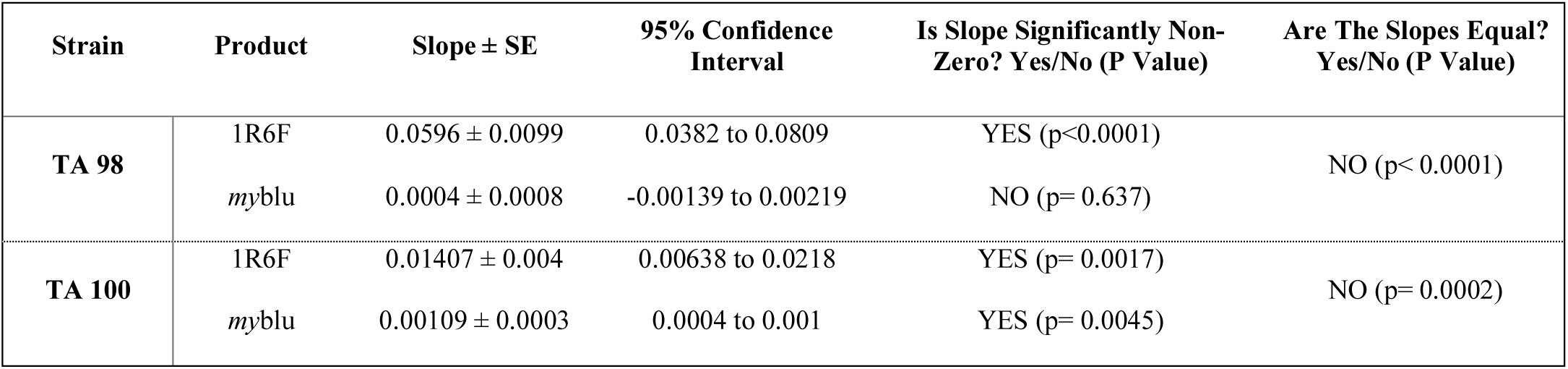
Linear regression analysis of mutagenic activity in *Salmonella tiphimurium* (TA98 and TA100) with S9 metabolic activation.

**Table 5.** Linear regression analysis of mutagenic activity in *Salmonella tiphimurium* (TA98 and TA100) without S9 metabolic activation.

| Strain | Product | Slope ± SE | 95% Confidence Interval | Is Slope Significantly Non-Zero? Yes/No (P Value) | Are The Slopes Equal? Yes/No (P Value) |
| --- | --- | --- | --- | --- | --- |
| TA 98 | 1R6F | 0.07815 ± 0.01045 | 0.0556 to 0.1007 | YES (p<0.0001) | NO (p< 0.0001) |
| | myblu | -0.0001 ± 0.001 | -0.003 to 0.002 | NO (p= 0.91) | |
| TA 100 | 1R6F | 1.102 ± 0.2120 | 0.6440 to 1.560 | YES (p= 0.0002) | NO (p< 0.0001) |
| | myblu | 0.00095 ± 0.0002 | 0.0005 to 0.0014 | YES (p= 0.0005) | |

**Figure 3.**
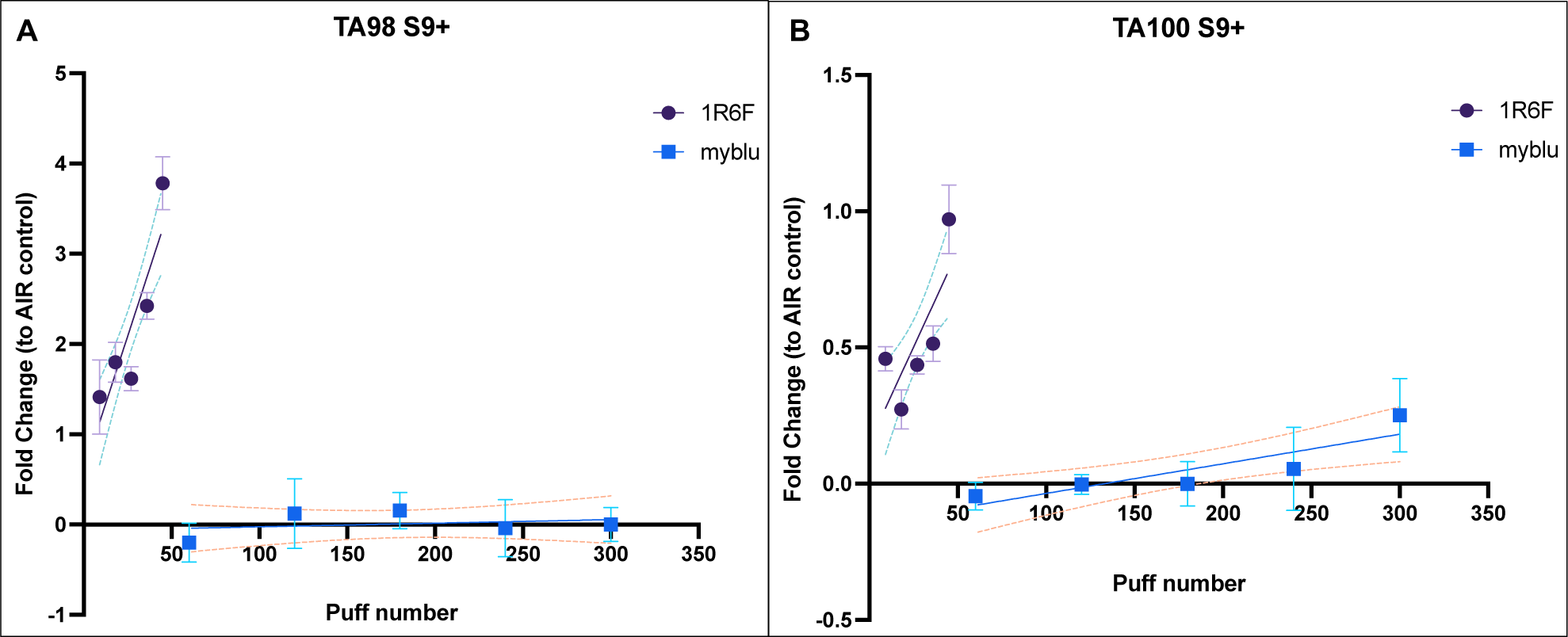
Mutagenicity evaluation by AMES test in *Salmonella typhimurium* TA98 (**A**) and TA100 (**B**) with S9 metabolic activation after exposure to 1R6F cigarette smoke (violet round dot) or myblu aerosol (blue square dot). Data are reported as Fold change to AIR control. Each data point represents the mean ± standard deviation (SD). The dashed lines represent the 95% confidence interval of the regression line.

Similar results were observed when the AMES assay was performed without S9 metabolic activation. Indeed, significant dose-dependent increase in revertants was observed in both TA98 (up to approximately 3.5-fold change to AIR control; p< 0.0001) and TA100 (up to 45 fold change to AIR control; p< 0.0001) without S9 metabolic activation after exposure to 1R6F cigarette smoke. The exposure to *my*blu aerosol did not induce any significant increase in revertants in both strains (Fig. 4). The linear regression results of mutagenic activity in *Salmonella tiphimurium* (TA98 and TA100) without S9 metabolic activation are reported in table 2.

**Figure 4.**
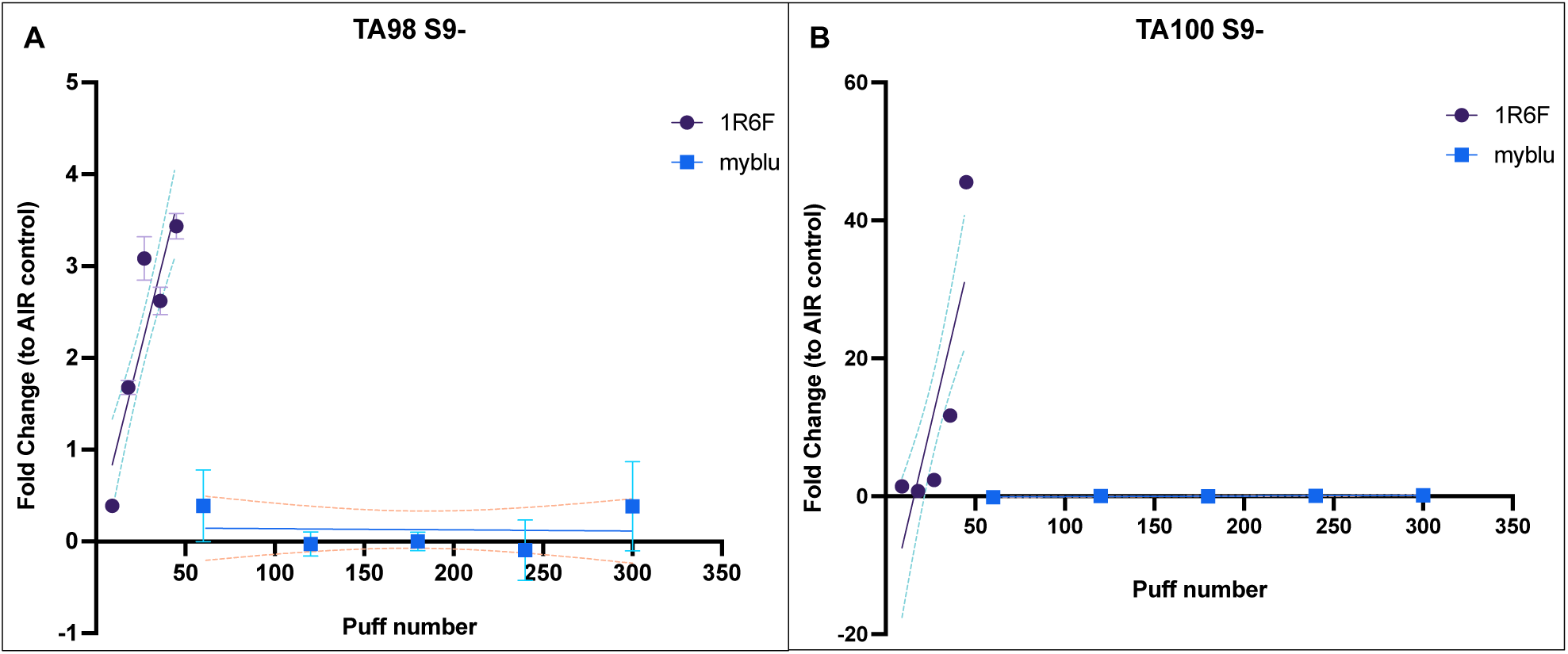
Mutagenicity evaluation by AMES test in *Salmonella typhimurium* TA98 (**A**) and TA100 (**B**) without S9 metabolic activation after exposure to 1R6F cigarette smoke (violet round dot) or *my*blu aerosol (blue square dot). Data are reported as Fold change to AIR control. Each data point represents the mean ± standard deviation (SD). The dashed lines represent the 95% confidence interval of the regression line.

### Genotoxicity effect of 1R6F cigarette smoke and *my*blu aerosol

Due to different exposure system, not able to perform smoke dilution, we performed a dose-response curve in order to establish the EC50 dose for the V79 cells exposed to 1R6F cigarette smoke: the calculated EC50 value for V79 cells was 3.149 puffs. We, then, performed the IVM assay with and without S9 metabolic activation after exposure from 1 to 4 puffs of 1R6F cigarette smoke. For *my*blu exposure the same puff numbers by Rudd et al. were used (20-100 puffs).

When the IVM assay was performed with the S9 metabolic activation, high cytotoxicity was observed for both 1R6F and *my*blu. Particularly, marked cytotoxicity was observed for V79 cells exposed to cigarette smoke to the point of being unable to perform micronuclei counts. The cause of this cytotoxicity was attributed to the S9 mix, which is known to be cytotoxic. Especially for the IVM assay with S9 of 1R6F, the cytotoxic effect of both S9 mix and undiluted cigarette smoke have added up, making the essay unfeasible. Whereas, we were able to perform micronuclei quantification in order to perform IVM assay for *my*blu despite the S9 cytotoxicity. The regression slope was not different from zero (p= 0.5) (Fig. 5), and all the micronuclei frequencies corresponding to each *my*blu puff number were not different from AIR control.

**Figure 5.**
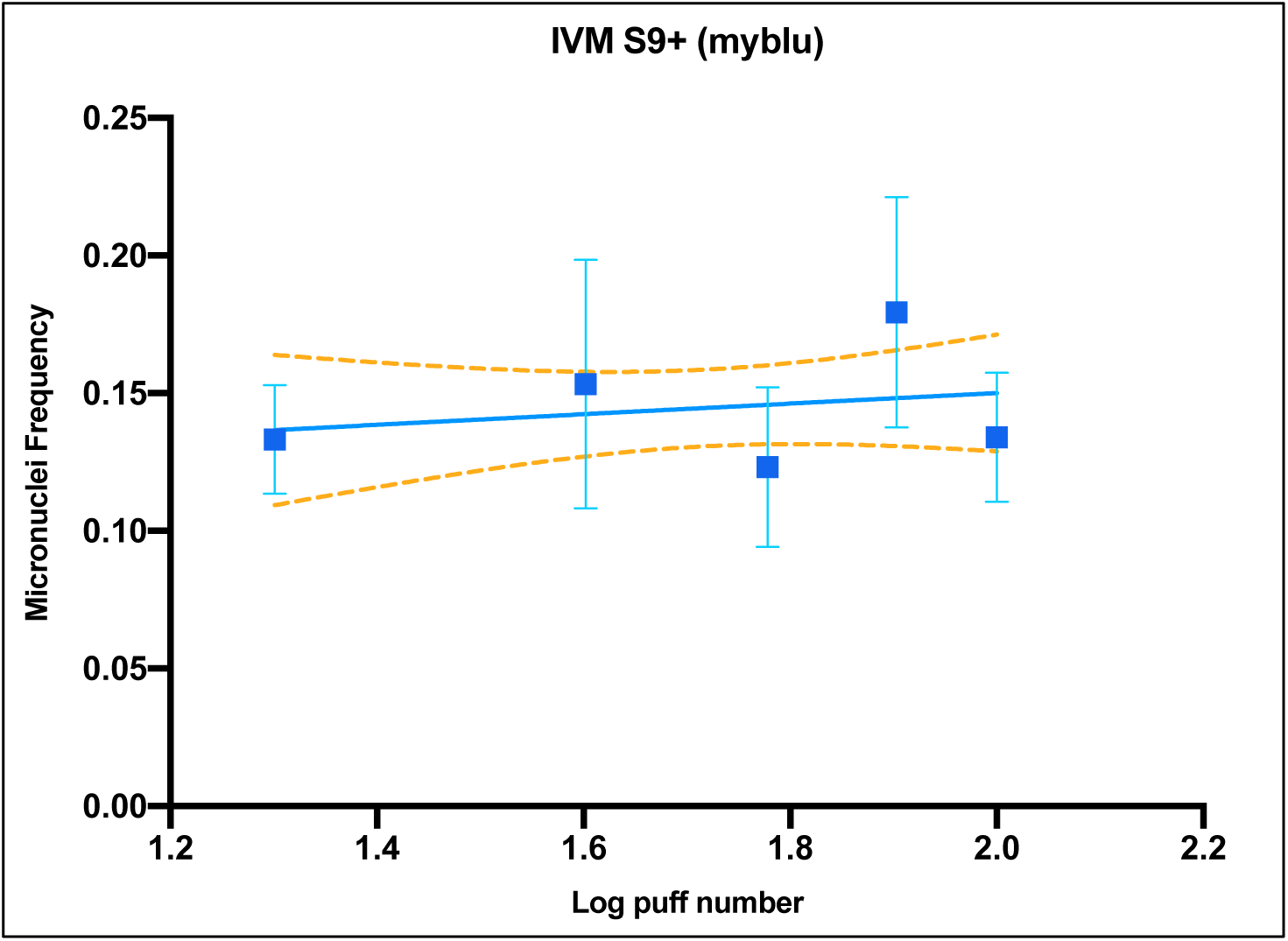
Genotoxicity evaluation by *in vitro* micronucleus assay with S9 activation in V79 cells after exposure to myblu aerosol (from 20 to 100 puffs) at the air liquid interface. Data are reported as micronuclei frequency. Each data point represents the mean ± standard deviation (SD). The dashed lines represent the 95% confidence interval of the regression line.

Because of the high cytotoxicity of S9 mixture and since the OECD n. 487 guideline reported that IVM assay can be performed with or without S9 activation, we performed this assay without S9 mixture in addition to what was done by Rudd and colleagues. The results of IVM assay without S9(S9-) are shown in figure 5. Increased micronuclei frequency was observed for the 1R6F exposure, but not in a dose-dependent manner due to high cytotoxicity of undiluted smoke (Fig. 6A). However, all the 1R6F puff numbers (from 1 to 4) induced significant increments of micronucleus frequency (p< 0.0001) compared to AIR control (Fig. 6B). Instead, no significant increase in micronucleus frequency was observed after exposure to *my*blu aerosol until 100 puffs. Moreover, no difference was shown among the three negative controls (INC, ALI and AIR) for both IVM with and without S9 activation. Instead, the positive controls, including cytochalasin A (for IVM S9+) and mitomycin C (for IVM S9-), were significantly increased compared to the respective AIR controls (Cyto A p< 0.0001; Mito C p= 0.002).

**Figure 6.**
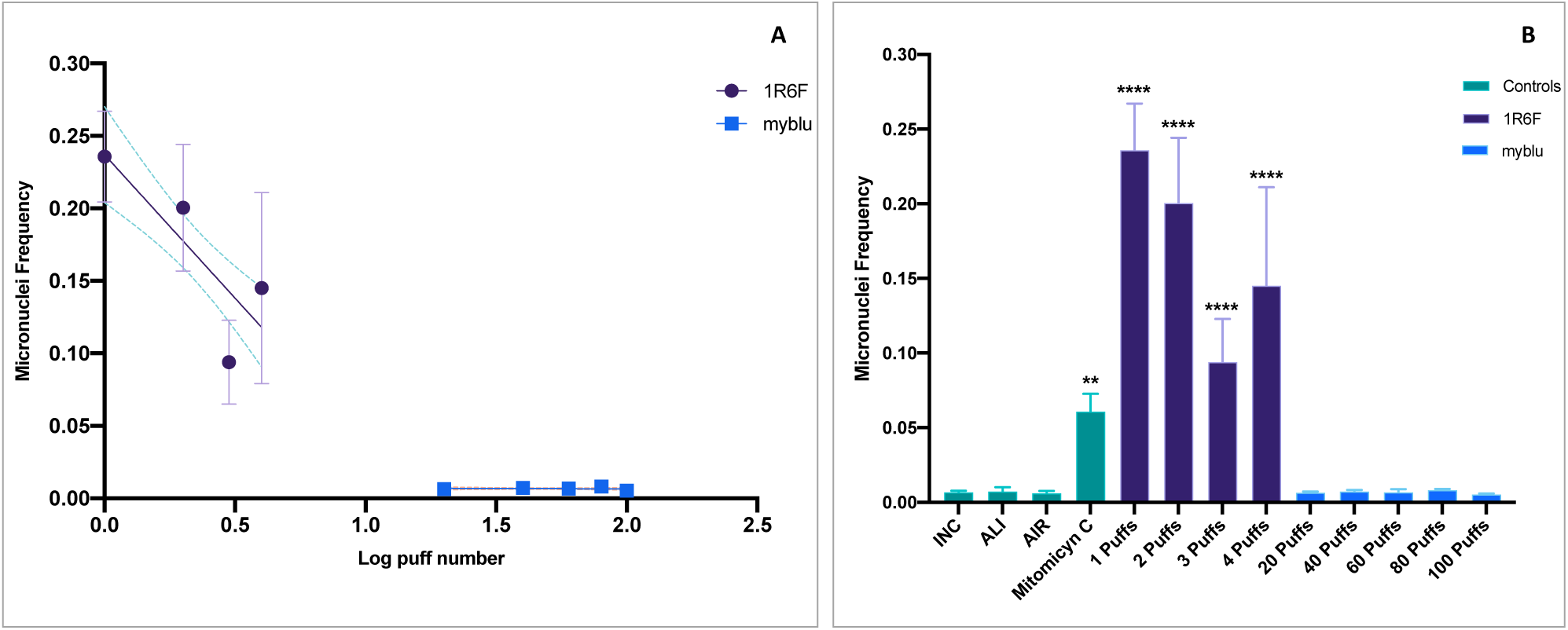
Genotoxicity evaluation by *in vitro* micronucleus assay without S9 activation in V79 cells after exposure to undiluted1R6F cigarette smoke or *my*blu aerosol at the air liquid interface. (**A**) Linear slopes of the dose–response IVM results for 1R6F (violet round dot) and *my*blu (blue square dot). The dashed lines represent the 95% confidence interval of the regression line. (**B**) Barplot representing IVM results, including both negative (INC; ALI; AIR) and positive (Mitomycin C) controls, 1R6F, and *my*blu results. All data are reported as micronuclei frequency. Each data point or bar represent the mean ± standard deviation (SD). ** p< 0.01; ****p< 0.0001 compared to AIR control.

## Discussion

This study replicated the work by Rudd and colleagues, which compared the *in vitro* toxicity of the *my*blu e-cigarette aerosol with those of cigarette smoke. They performed a standard toxicological battery of three assays used for product assessment and regulatory applications: the NRU assay to assess cytotoxicity [15], the bacterial reverse mutation (Ames) assay to evaluate mutagenicity [18], and the *in vitro* micronucleus assay to measure genotoxicity [19]. Their results indicated that e-cigarette aerosol was low cytotoxic, and it did not show any mutagenic or genotoxic activity unlike the 3R4F cigarette smoke, which showed high cytotoxic, mutagenic and genotoxic activity.

Despite some different methodological aspects in our study, we obtained results similar to those obtained by Rudd and colleagues. The main methodological differences were: (i) the use of 1R6F reference cigarette in place of the 3R4F reference cigarette because the latter is no longer produced by the Center for Tobacco Reference Products (University of Kentucky); (ii) the different smoking and vaping apparatus. Our laboratory is equipped with separate machines, the LM1 smoking machine and the LM4E vaping machine (Borgwaldt, Hamburg, Germany). Instead, Rudd and colleagues used the SAEIVS five-port smoking/vaping machine for generation of both smoke and aerosol. Also, the SAEVIS machine is able to perform dilution of smoke/aerosol towards the LM1 and LM4E machine that were not designed to perform dilutions; (iii) the different smoke/aerosol ALI exposure *in vitro* system. The smoke/aerosol exposure *in vitro* system (SAEIVS) used by Rudd and colleagues was designed to expose cells in 96 and 24 multi-well plates, only the latter with transwell inserts. Instead, our *in vitro* ALI exposure system (described in the “methods” section) allows the cell exposure with transwell inserts of all diameters by the use of a dedicated exposure chamber. All these differences have been filled by implementing some modifications to the protocols used in the original work as described in the “methods” section.

NRU assay was performed both in LAB-A and in LAB-B. Our results confirmed the higher cytotoxity of 1R6F cigarette smoke compared to the e-cigarette aerosol as showed by Rudd and colleagues. However, the calculated EC50 for the 1R6F smoke (1.71 puffs) was different from that obtained in the original work (0.236 puffs). In addition, we did not observe the same cytotoxic effect for the *my*blu aerosol. Indeed, the low cytotoxicity induced by myblu aerosol did not allow us to calculate the value of EC50. But we observed only a reduced cell viability, around the 80% of viability, starting from the 80 puffs to 140 puffs. These differences in results may be ascribed to the different ALI exposure apparatus. Rudd and colleagues exposed BEAS-2B cells seeded in the 96-well plate, but the cells do not have medium in the basal face of cells by this type of exposure. Then it is not a real ALI exposure because the cells are dry as the apical medium is taken to perform an air-interface exposure. As a result, part of the cytotoxicity observed by Rudd and colleagues could be due to conditions that are not optimal for normal cell health. Conversely, we exposed BEAS-2B cells using transwell inserts placed into the exposure chamber filled with culture medium at the basal compartment that provides nutrition for cells through the transwell membrane. This exposure apparatus provides the optimal environment to avoid the cells to dry out, especially when performing long exposures (10 to 77 minutes, in this case). The same ALI-exposure system combined with cytotoxicity evaluation was successfully used in our previous works [14, 21-23], and other published works [24, 25]. Moreover, several *in vitro* toxicity studies used the ALI exposure with cell cultures by using appropriate apparatus developed by dedicated manufactures, such as the VITROCELL® and CULTEX® system, that simulate real *in vivo* exposure conditions [26, 27].

The AMES test was performed only in the LAB-A, as reported by Rudd and colleagues, with *Salmonella typhimurium* TA98 and TA100 strains, which are particularly relevant for tobacco products since they have been shown to be sensitive to combustion products [28, 29]. Unlike the original work, we conducted the AMES test with and without S9 metabolic activation. Our results showed that neither TA98 nor TA100 with and without S9 showed a mutagenic response after myblu aerosol exposure even at high doses (from 60 to 300 puffs), as opposed to what has been observed for 1R6F cigarette smoke with minor doses (from 9 to 45 puffs). These results are aligned with what Rudd and colleagues showed in their work, even though we used different exposure machines. The mutagenicity evaluation of e-cigarette aerosol by AMES assay has been also reported in literature with similar results to those reported in this work [9, 30-32].

Genotoxicity evaluation was conducted by LAB-A in a similar way to those reported by Rudd and colleagues, with the following exceptions: (i) we added the genotoxicity evaluation without S9 metabolic activation to improve their results; (ii) the exposure of 1R6F cigarette smoke was performed undiluted. Though, we experienced some methodological issues following their protocol. Indeed, they reported the use of S9 mix at 10%, but using the same concentration we observed a massive cell mortality (more than 50%) especially for 1R6F cigarette smoke that affect the observation of micronuclei in V79 cells. Instead, we were able to perform the micronuclei evaluation with S9 for the V79 cells exposed to myblu aerosol, showing no genotoxic effect. Based on literature data, we found that the S9 mix has a full set of liver metabolic enzymes, but it displays high cytotoxicity in cell-based assays [33]. Consequently, we performed a dose-response curve with different concentrations of S9 mix (from 1% to 5%), and we observed that cell viability decreased with the increment of S9 enzymatic mix percentage. Probably, the high cytotoxicity levels observed in the IVM assay with S9 are due to the sum of undiluted whole smoke cytotoxicity plus the S9 enzymatic mix cytotoxicity. A limitation of IVM assay is that higher cytotoxicity levels may induce chromosome damage as a secondary effect of cytotoxicity, then it is suggested not to exceed 50% cytotoxicity [19]. Indeed, the IVM assay without the S9 metabolic activation allow us the micronuclei count for both 1R6F cigarette smoke and myblu aerosol. High genotoxicity (higher than positive control) was showed for the 1R6F cigarette smoke, although we did not observe a clear dose response. No genotoxicity was observed for all the *my*blu exposure conditions. In line with our results there is the work by Thorne et al., which showed the highest responsivity of V79 cells to cigarette smoke constituents after an extended recovery period without S9 [10].

## Conclusions

In conclusion, our findings confirmed the results on low toxicity profile of *my*blu e-cigarette obtained by Rudd and colleagues, despite some differences in methodology. Moreover, our study covered some methodological gaps and limitations in the original work, including the non-optimal ALI exposure for the cytotoxicity evaluation and improved mutagenicity and genotoxicity results by adding experiments without S9 metabolic activation as recommended in the OECD guidelines. Overall, this replication study supports the tobacco harm reduction strategy as having the potential to substantially reduce exposure to toxic combustion agents in adult smokers. Future studies are needed to advance *in vitro* methods in order to evaluate the long-term effects of electronic nicotine delivery systems.

